# Multivalent Clustering of Adhesion Ligands in Nanofiber-Nanoparticle Composites

**DOI:** 10.1101/2020.06.12.148288

**Authors:** Dounia Dems, Ronit Freeman, Thibaud Coradin, Samuel I. Stupp, Carole Aimé

**Author notes:** These authors contributed equally to the work (Dounia Dems, Ronit Freeman). Ecole Normale Supérieure, CNRS-ENS-UPMC UMR 8640, 24 Rue Lhomond, Paris, 75005, France.

## Abstract

Because the positioning and clustering of biomolecules within the extracellular matrix dictates cell behaviors, the engineering of biomaterials incorporating bioactive epitopes with spatial organization tunable at the nanoscale is of primary importance. Here we used a highly modular composite approach combining peptide amphiphile (PA) nanofibers and silica nanoparticles, which are both easily functionalized with one or several bioactive signals. We show that the surface of silica nanoparticles allows the clustering of RGDS bioactive signals leading to improved adhesion and spreading of fibroblast cells on composite hydrogels at an epitope concentration much lower than in PA-only based matrices. Most importantly, by combining the two integrin-binding sequences RGDS and PHSRN on nanoparticle surfaces, we improved cell adhesion on the PA nanofiber/particle composite hydrogels, which is attributed to synergistic interactions known to be effective only for peptide intermolecular distance of *ca*. 5 nm. Such composites with soft and hard nanostructures offer a strategy for the design of advanced scaffolds to display multiple signals and control cell behavior.

## 1. INTRODUCTION

The natural extracellular matrix (ECM) surrounding cells plays a critical role in directing cell function by providing essential structural and biochemical cues. One mechanism by which ECMs regulate cell signaling is clustering of biological ligands with variable densities and separation.^1–3^ For example, focal adhesions are triggered by the formation of an effective integrin cluster with a specific lateral spacing. This has been experimentally demonstrated by controlling the density and interspacing of arginine-glycine-aspartate (RGD) ligands in synthetic materials, showing that the peptide spacing (*ca.* 70 nm) within a local cluster is more essential than its bulk density to trigger cell adhesion.^4–7^ In addition to RGD clustering, integrin-binding proteins contain domains that operate synergistically with RGD to elicit cell response. For instance, the PHSRN sequence within fibronectin synergizes with RGD in a distance-dependent manner.^8–9^ Different approaches have been developed to control ligand positioning and inter-ligand distances, including their conjugation onto amphiphilic^10–13^ and PEGylated constructs,^14,15^ oligopeptide backbones,^16–18^ DNA constructs^19–21^ or the functionalization of titanium surfaces to display distinct bioactive motifs in a chemically-controlled fashion.^22^ The use of self-assembling peptide amphiphiles (PAs), which consist of a short peptide sequence linked to a hydrophobic alkyl tail, has been particularly promising in engineering bioactive artificial scaffolds for cells.^23,24^ The facile incorporation of multiple bioactive signals at controlled concentrations, together with their structural similarity to extracellular matrix fibres makes PA assemblies useful as bioactive artificial extracellular matrix components for cell signalling.^25,26^ Interestingly, peptide amphiphile supramolecular systems were shown to have both fully dynamic and kinetically inactive areas in the aggregate, which can be used to generate useful cluster morphologies.^27,28^

Over the last few years, the nanocomposite approach has emerged as an efficient alternative to generate biofunctional scaffolds.^29^ Bionanocomposites based on the association between bio-based polymers and inorganic colloids combine the chemical diversity, hierarchical structure and biocompatibility of biomacromolecules with the robustness and functionality of the inorganic phase.^30^ Depending on the chemical nature of the nanoparticles (NPs), different properties can be imparted to the resulting composite to design conductive, optical and magnetic devices, and also to tune the mechanical properties and the bioactivity of hydrogels.^31,32^ Silver NPs have often been incorporated within matrices of biological or synthetic origin for their antimicrobial properties,^33–36^ as well as to design plasmonic sensors^37^. Gold,^38–40^ cobalt,^41^ nickel^42^ and copper^43,44^ metal NPs together with iron oxides NPs^45–48^ have also been encapsulated to design composites, many of which finding applications in drug delivery. In parallel, the incorporation of silica nanoparticles (SiNPs) has been shown to enhance the mechanical properties of hydrogels,^49–53^ enhances biological activity of biomaterials^54^ and has been widely studied in the field of drug delivery.^55–60^ SiNPs are particularly interesting candidates due to their low cost, limited cytotoxicity, ease of synthesis, and the versatility of sol-gel chemistry that offers various routes to conjugate biomolecules at the NP surface, while preserving their molecular recognition properties.^61,62^

Here we combine self-assembled PA matrices with SiNPs to design novel SiNP-PA composite biomaterials (**Figure 1**). The ability to independently modify the chemistries of both PA and NP substrates to link distinct bioactive motifs on which cells would grow allows us to cluster signals in variable patterns positioned through the composite material to impart biological functionality. We show that clustering of the fibronectin derived RGDS peptide on the surface of Stöber SiNPs (*ca*. 200 nm in diameter, **Figure 1A**) triggers cell adhesion onto SiNP-PA scaffolds. In addition, the multiple display of RGDS and PHSRN bioactive epitopes can be achieved within SiNP-PA composites (**Figure 1B**) to trigger synergistic effects on cell behavior. This strategy offers a unique modularity by the ability to introduce functionality through both the nanofiber scaffold and the incorporated modified NPs, making composites highly promising biomaterials to display bioactive sequences with synergistic effects.

**Figure 1.**
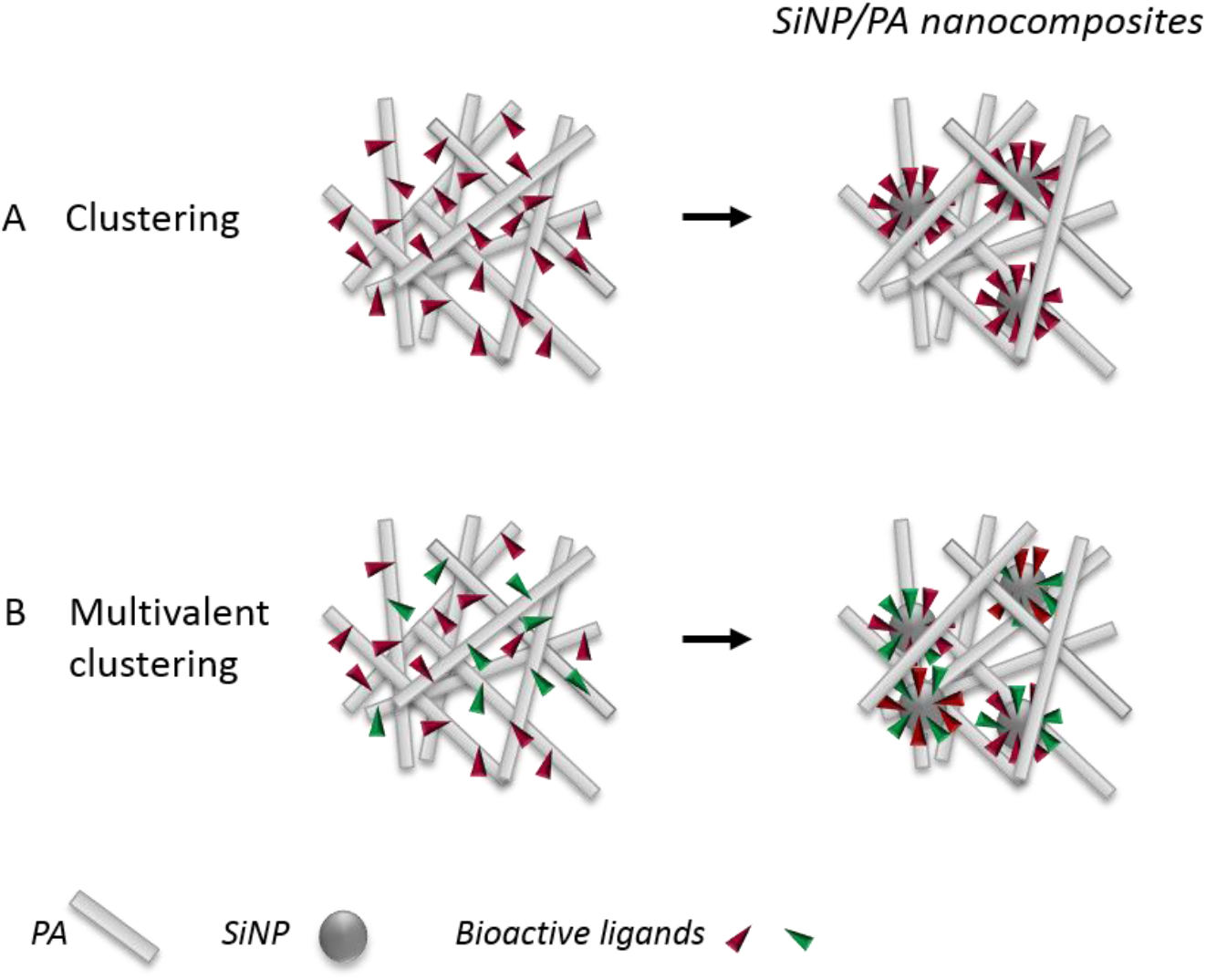
Schematic representation of the different composites. (A) Single ligand display in PA nanofibers (left) and clustered at the surface of SiNPs embedded in a PA matrix (right). (B) Simultaneous display of two bioactive signals in PA nanofibers (left) or after the clustering of two signals at the surface of SiNPs embedded in a PA matrix (right).

## 2. EXPERIMENTAL SECTION

### PA and Peptide synthesis

PAs and peptides were synthesized using a standard fluorenylmethyloxycarbonyl (Fmoc) solid phase peptide synthesis (SPPS) on Rink Amide MBHA resin as described previously.^63^ Amino acid couplings were performed either manually or on a CEM Liberty microwave-assisted peptide synthesizer. Rink Amide MBHA resin, Fmoc-protected amino acids and 2-(1H-benzotriazole-1-yl)-1,1,3,3-tetramethyluronium hexafluorophosphate (HBTU) were purchased from Novabiochem; Fmoc-NH-PEG_4_-CH_2_COOH was purchased from ChemPep Inc.; palmitic acid was purchased from Acros Organics; Fmoc-(4-amino)benzoic acid and Fmoc-(4-aminomethyl)benzoic acid were purchased from VWR and Chem-Impex International Inc., respectively. All other reagents and solvents were purchased from Sigma Aldrich and used as received. Fmoc deprotection was performed using 30% piperidine in N,N-dimethylformamide (DMF) and amino acid and palmitic acid couplings were performed with 4 molar equivalent (eq.) protected amino acid or palmitic acid, 3.95 eq. HBTU, and 6 eq. of *N*,*N*-diisopropylethylamine (DIEA) in DMF alone or in a solvent mixture of 1:1:1 DMF/dichloromethane (DCM)/*N*-methyl-2-pyrrolidone (NMP). The coupling reaction for the PEGylated amino acid was performed similarly to other standard Fmoc-protected amino acids, using Fmoc-PEGylated amino acid (3 eq.), HBTU (2.95 eq.), and DIEA (4.5 eq.) in DMF. For the coupling of Fmoc-(4-amino)benzoic acid and Fmoc-(4-aminomethyl)benzoic acid, both were converted into acid chloride first (procedure described below) to increase the coupling yields. The coupling reaction was performed by using 4 eq. of Fmoc-(4-amino)-benzoyl chloride or Fmoc-(4-aminomethyl)benzoyl chloride and 6 eq. of DIEA in NMP. Synthesized PA and peptide molecules were cleaved from the resin using a mixture of 95% trifluoroacetic acid (TFA), 2.5% water, and 2.5% triisopropylsilane (TIPS). After removing TFA by rotary evaporation, the product was precipitated with cold diethyl ether, dried, and purified using preparative scale reverse phase high performance liquid chromatography on a Varian Prostar Model 210 system equipped with a Phenomenex Jupiter Proteo column (C12 stationary phase, 10 mm, 4 μm particle size and 90 Å pore size, 150 × 30 mm). A linear gradient of acetonitrile (2 to 100%) and water with 0.1% ammonium hydroxide (added to aid PA solubility) was used as the mobile phase for purification. Electrospray ionization mass spectrometry (Agilent 6510 Q-TOF LC/MS) was used to identify the pure fractions (Figure S1,S2), which were then combined together and lyophilized after removing excess acetonitrile by rotary evaporation.

### Synthesis of silica particles

Silica particles (*ca*. 200 nm in diameter) were synthesized by the Stöber process using 32 mL ultrapure water, 600 mL absolute ethanol (VWR, GPR RectaPur), 45 mL ammonium hydroxide solution (25%, Carlo Erba), and 21 mL tetraethyl orthosilicate (TEOS 98%, Aldrich).^64^ After extensive washing, SiNPs were characterized by TEM and DLS (Figure S3).

### Synthesis of peptide-conjugated SiNPs

The synthesis of peptide-conjugated SiNPs proceeded in three steps, whose success was checked by zeta potential ζ measurements (Figure S4).

#### Amine functionalization of SiNPs

Stöber particles were first functionalized with amine groups with (3-Aminopropyl)triethoxysilane (APTES, 99%, Aldrich). Typically, 0.77 g of silica particles were redispersed in a mixture of 76.6 mL ethanol and 1.7 mL ammonium hydroxide solution before addition of 0.75 ml APTES (4.2 mmol.g^−1^ silica). The mixture was stirred for 18 h at RT. Subsequently, the reaction mixture was heated to 80°C and the total volume was reduced to approximately two-third by distillation of ethanol and ammonia at ambient pressure. The mixture was left to cool down to RT and was subsequently washed three times with ethanol (by centrifugation at 12 000 rpm for 15 min) before drying under vacuum. Successful surface modification was ascertained by the increase of the ζ potential value of the recovered nanoparticles at all pH values compared to initial SiNPs.

#### Dibenzocyclooctyne-N-hydroxysuccinimidyl ester grafting on SiNP-APTES

Amine-bearing silica nanoparticles were redispersed in a phosphate buffer solution at pH 8.3 before addition of 94 μmol of Dibenzocyclooctyne-N-hydroxysuccinimidyl ester (DBCO-NHS, Aldrich) in Dimethylsulfoxide (DMSO, Aldrich) (3 mmol.g^−1^ silica). The mixture was stirred for 12 h at RT and subsequently washed three times with water (by centrifugation at 12 000 rpm for 15 min) before drying under vacuum. The recovered nanoparticles exhibited ζ potential values very similar to the initial SiNPs, indicating the successful coupling of the surface amine groups with DBCO.

#### Peptide grafting via Click Chemistry

SiNP-DBCO were redispersed in water before addition of 1.2 μmol of the azide-bearing peptide (RGDS, PHSRN, RGES) in DMSO (4 mmol.g^−1^ silica). The mixture was stirred for 12 h at RT and subsequently washed three times with water (by centrifugation at 12 000 rpm for 15 min). The measured variations of the ζ potential values with pH were in agreement with the main ionizable groups of the amino acid sequence of each peptide (guanidine, imidazole for PHSRN; guanidine, carboxylate for RGDS and RGES).

### Quantification of surface functionalization using Cy3-Azide

SiNP-DBCO or nude SiNPs were redispersed in water before addition of 1.2 μmol of Cy3-azide (Cy3-N3, 90%, Aldrich) in DMSO (4 mmol.g^−1^ silica). The mixture was stirred for 12 h at RT and subsequently washed as many times as necessary (at least 5 times) with water (by centrifugation at 12 000 rpm for 15 min). Absorbance and fluorescence of the samples were then measured to quantify the conjugation rate at the surface of SiNPs, providing a density of 0.2 Cy3 per nm^2^ of silica surface (Figure S5).

### Peptide Amphiphile (PA) and SiNP/PA Composite Gel Preparation

The desired amount of PA powder was weighed out in an Eppendorf tube in order to make 100 μL of a 1 wt% PA stock solution in H_2_O. The PA solution was subsequently annealed at 80°C in a PCR machine for 30 min and slowly cooled down to room temperature (RT) over 90 min. The self-assembled structures resulting from the different PA mixtures were characterized using TEM (Figure S6). SiNP/PA composite gels were prepared following the same protocol except that a water suspension of SiNP was added to the PA solution at different ratios and the mixture was sonicated before annealing.

### Rheology

Rheological measurements were performed on a Paar Physica MCR 300 oscillating plate rheometer equipped with a 25 mm diameter cone-plate geometry and a gap of 0.05 mm. PA or PA+SiNP solutions at a PA concentration of 0.5% (w/v) in water were pipetted (180 μL) onto the rheometer plate and gelled by exposure to 50 μL CaCl_2_ solution (20 mM CaCl_2_, 150 mM NaCl). All measurements were done at 25°C, and the gels were allowed to equilibrate for 5 min at 0.1% strain prior to measurement. Data were collected at 0.1% strain over a frequency range of 1 to 100 s^−1^ and all measurements repeated 3 times.

### Transmission Electron Microscopy

PA and SiNP/PA samples were deposited and dried on 300 square mesh carbon-coated copper grids (Ted Pella, Redding, CA) and stained with 0.5% uranyl acetate (UA) solution. Images were obtained using a Hitachi HT-7700 Biological TEM (Hitachi High Technologies America, Schaumburg, IL) equipped with a LaB6 filament working at an accelerating voltage of 100 kV.

### Preparation and imaging of SiNP/PA layers

**SiNP/PA** layers were prepared on sterile glass coverslips (12 mm diameter) or tissue culture plates. The sample surface was first coated with 0.01% (w/v) poly-D-lysine (Aldrich) in milliQ water. A 1% (w/v) PA solution in milliQ water supplemented with SiNPs at various ratios was added onto the surface and the layer was gelled with a 10 mM CaCl_2_ aqueous solution. These layers were characterized by SEM (Figure S7). SiNP-Cy3/PA layers were also prepared. The PA were stained by DAPI and images of the sample were obtained using an inverted confocal laser scanning microscope (Nikon A1R).

### Cell culture

NIH 3T3 mouse embryonic fibroblasts were maintained in growth medium containing Dulbecco’s Modified Eagle’s Medium (DMEM) with high glucose, supplemented with 10% fetal bovine serum (FBS) and 1% penicillin-streptomycin (P/S). The cells were grown in 75 mm^2^ flasks (BD Falcon) and passaged every three days. All culture reagents were purchased from Gibco. For cell morphology experiments on PA layers, fibroblasts were seeded at a low density (5,000 cells per well) in order to minimize cell–cell contacts, and incubated (at 37°C, 5% CO_2_) under serum free condition (DMEM + 1% P/S). The serum free media was used to eliminate any interference from serum adsorption to the nanofibers. Within the time-period of experiment (4h30), no adverse cellular responses were observed from serum deprivation or serum shock after a transfer from serum containing growth media.

### Confocal microscopy

Cells were fixed with 4% paraformaldehyde in PBS and 1 mM CaCl_2_ for 30 min at RT. For immunostaining, fixed samples were first permeabilized with 0.1% Triton X-100 in PBS (5 min, RT). Actin filaments were fluorescently labeled with AlexaFluor-488-conjugated phalloidin (Life Technologies; 1: 200 dilution, 1 h at RT) for visualization. Cell nuclei were counterstained with DAPI (Life Technologies).Images of fluorescently stained samples were obtained using an inverted confocal laser scanning microscope (Nikon A1R) or TissueGnostics cell imaging and analysis system mounted to an upright microscope (Zeiss). Cell morphology was quantified from phalloidin stained fluorescent images acquired by a 20× objective from randomly selected regions on the coverslip. Acquired grayscale images were background subtracted and thresholded to convert into binary images using ImageJ software (NIH).

### Scanning electron microscopy (SEM)

Cells on PA or SiNP/PA-coated glass coverslip were fixed with 2.5% glutaraldehyde in PBS (containing 1 mM CaCl_2_) for 1 hour at RT. Fixed samples were dehydrated by exposure to a graded series of water-ethanol mixture. Once in 100% ethanol, samples were dried at the critical point of CO_2_ using a critical point dryer (Tousimis Samdri-795) to preserve structural details. Dried samples were then coated with 14 nm of osmium using an osmium plasma coater (Filgen, OPC-60A), and imaged using a Hitachi S-4800 Field Emission Scanning Electron Microscope working at an accelerating voltage of 5 kV.

### Statistical Analysis

Statistical analysis was performed using Graphpad Prism v.6 software. Analysis of variance (ANOVA) with the Turkey’s Multiple Comparison test was used for all multiple group experiments. P values < 0.05 were deemed significant. Values in graphs are the mean and standard error of mean.

## 3. RESULTS

### 3.1. Preparation and characterization of SiNP-PA composites

C_16_V_3_A_3_E_3_ (**Figure 2**) is a PA previously shown to form nanofiber networks when intermolecular electrostatic repulsive interactions are screened by a salt solution.^28^ This PA was biofunctionalized by conjugating the peptide epitopes RGDS (arginine, glycine, aspartic acid, serine) and PHSRN (proline, histidine, serine, arginine, asparagine) leading to the two conjugates PA-RGDS (C_16_V_3_A_3_E_3_-G_5_-RGDS) and PA-PHSRN (C_16_V_3_A_3_E_3_-G_5_-PHSRN) (**Figure 2**).

**Figure 2.**
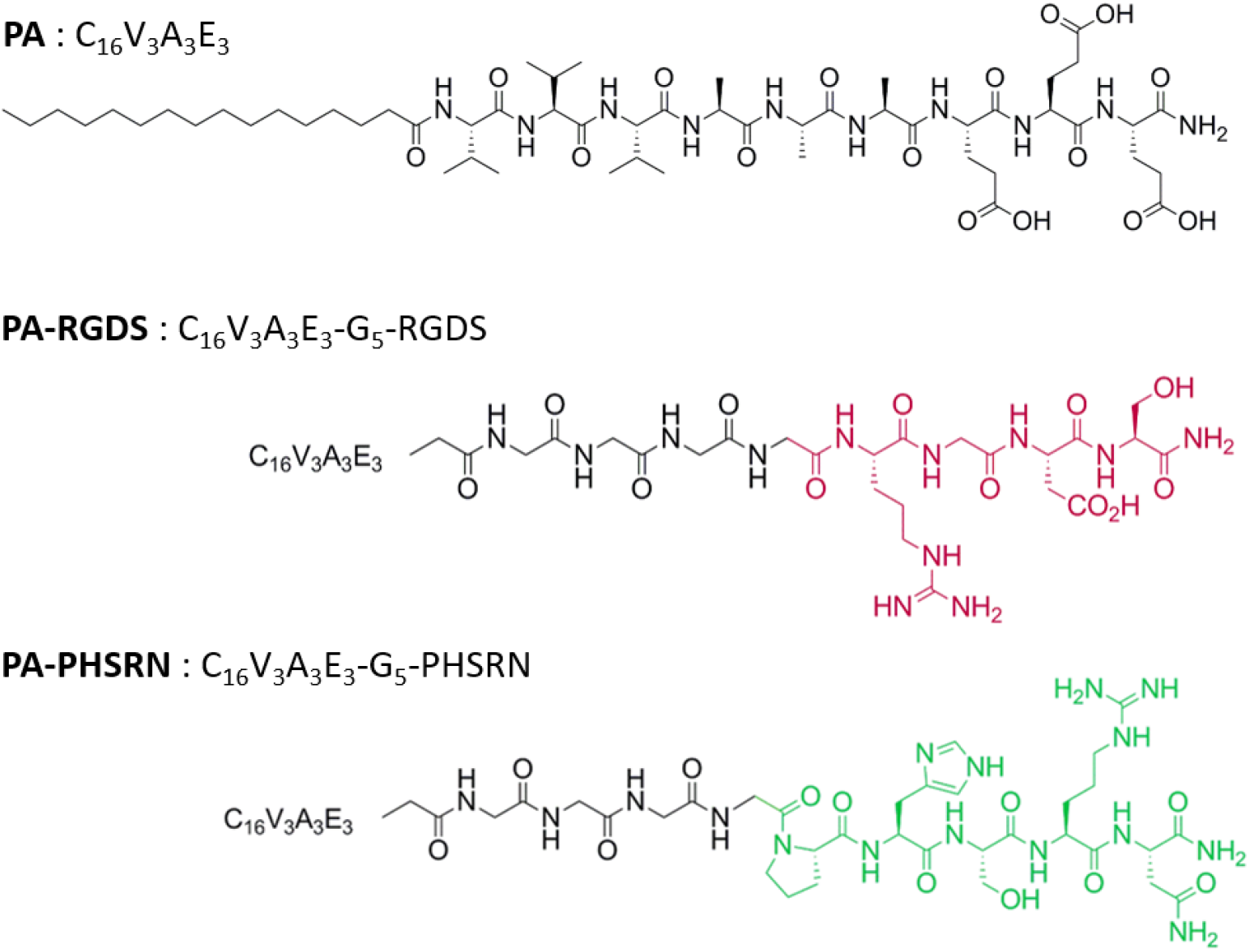
Molecular structure of the C_16_V_3_A_3_E_3_base PA and of the two epitope-conjugated PAs (PA-RGDS and PA-PHSRN). A biofunctional matrix can be obtained upon mixing the base PA with the bioactive derivatives PA-RGDS and/or PA-PHSRN. No significant variation in the gel structure is observed up to 2.6 mol% of peptide epitope (**Figure S6**). The nanocomposite counterpart can be prepared by the incorporation of SiNPs functionalized with peptide epitopes that are conjugated to amine-modified SiNPs.

First, amine-modified SiNPs were mixed with a 1 wt% (10 mg.mL^−1^) PA solution before gel formation. In this case, the SiNP concentration (from 3 to 25 mg.mL^−1^) was selected so as to target a peptide epitope concentration of 0.2 to 2.6 mol%, assuming that all surface amines are modified with peptide epitopes. The different co-assemblies all formed a gel, incorporating SiNPs within the nanofiber network, as observed by TEM and SEM for 1.3 and 2.6 mol% SiNPs (**Figure 3A,B and D,E**). This indicates that the presence of SiNPs does not disturb the PA self-assembly. The localization of SiNPs within the PA matrix could further be visualized by the conjugation of azide-cyanine 3 dye to SiNPs and of DAPI to the PA. Observations by confocal microscopy confirmed the good dispersion of SiNPs within the 3D gel (**Figure 3C,F**).

**Figure 3.**
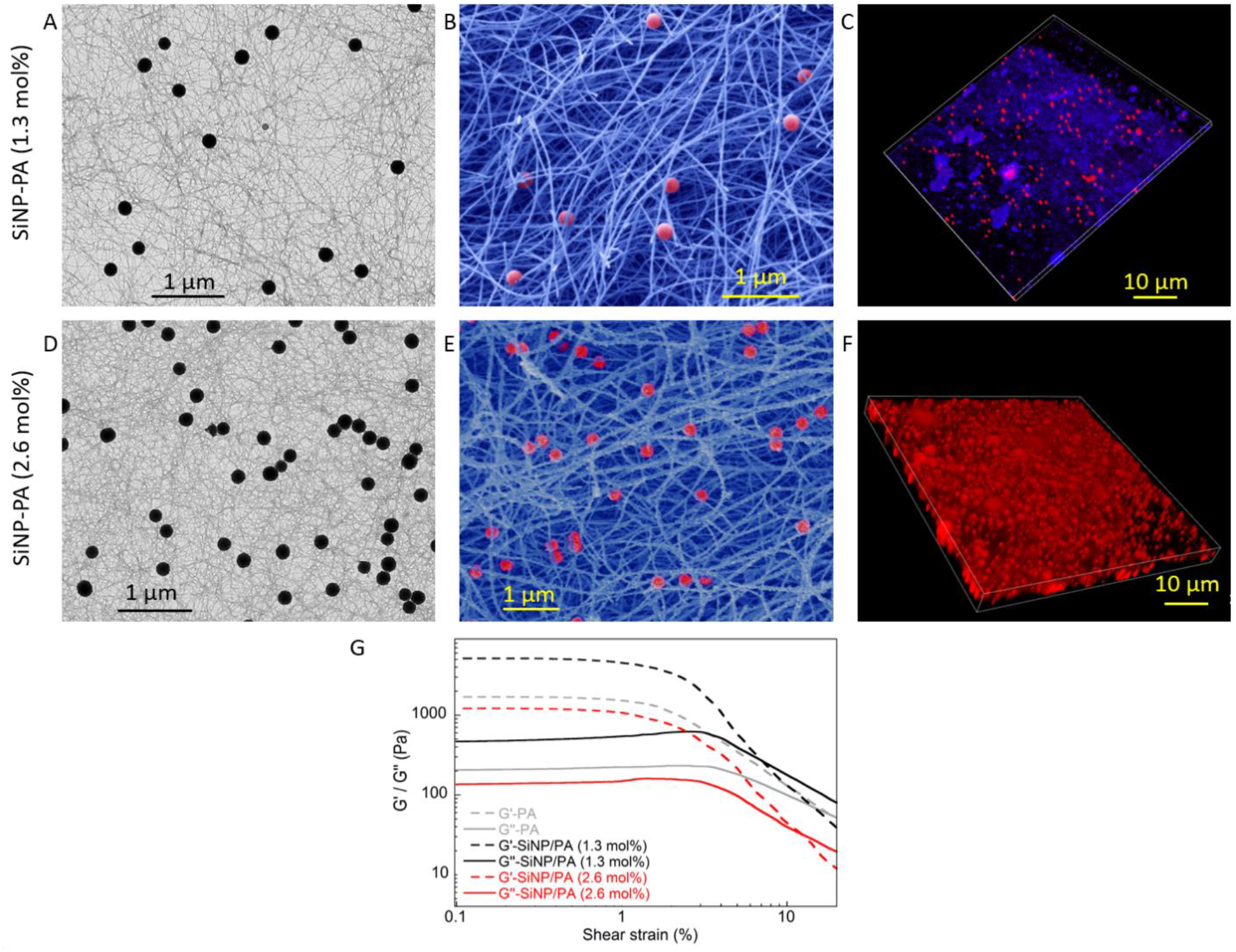
TEM, SEM (colored image) and confocal images (red: Cy3; blue:DAPI) of SiNP/PA at (A-C) 1.3 mol% and (D-F) 2.6 mol% SiNPs. (G) Rheology measurements of the PA alone and of SiNP/PA at 1.3 and 2.6 mol%.

Next, the rheological properties of the PA and SiNP-PA (1.3 and 2.6 mol%) scaffolds were assessed, **Figure 3G**. The SiNP PA scaffolds remained in similar range of mechanical stability with storage moduli in the range of 100-500 Pa.

### 3.2. Bioactivity of the single-peptide composite: the clustering effect

3T3 fibroblasts were cultured on PA-RGDS and SiNP-RGDS PA at different RGDS concentrations. Immunostaining for actin filament (phalloidin in green) and nuclei (DAPI in blue) showed low number of cells and no significant spreading of the cells in the absence of RGDS (**Figure 4Aa**). In contrast, improved cell adhesion and spreading was clearly observed for PA-RGDS matrices when reaching 2.6 mol% RGDS, and the cells showed formation of focal adhesions. The lower concentrations (0.6 and 1.3 mol%) of PA-RGDS were not sufficient to promote cell adhesion and spreading (**Figure 4Ab-c**). Interestingly, the incorporation of SiNP-RGDS showed a positive effect on cell spreading at a concentration as low as 0.6 mol% (Figure 4Af). Quantitative assessments obtained by image analysis confirmed that PA-RGDS at 2.6 mol% and SiNP-RGDS PA at 0.6 mol% were equally efficient in promoting cell spreading. .Incorporation of SiNPs bearing the mutated peptide RGES (**Figure 4Ai,j**) or non-functionalized SiNPs (Figure S8) did not result in cell spreading. While it is difficult to completely rule out a local mechanobiology effect, the observed decrease in RGDS concentration required to improve cell spreading between the PA-RGDS and PA-NP composite systems suggests that differences in epitope display within the scaffold and the local high concentration of SiNP RGDS PA are playing a role.

**Figure 4.**
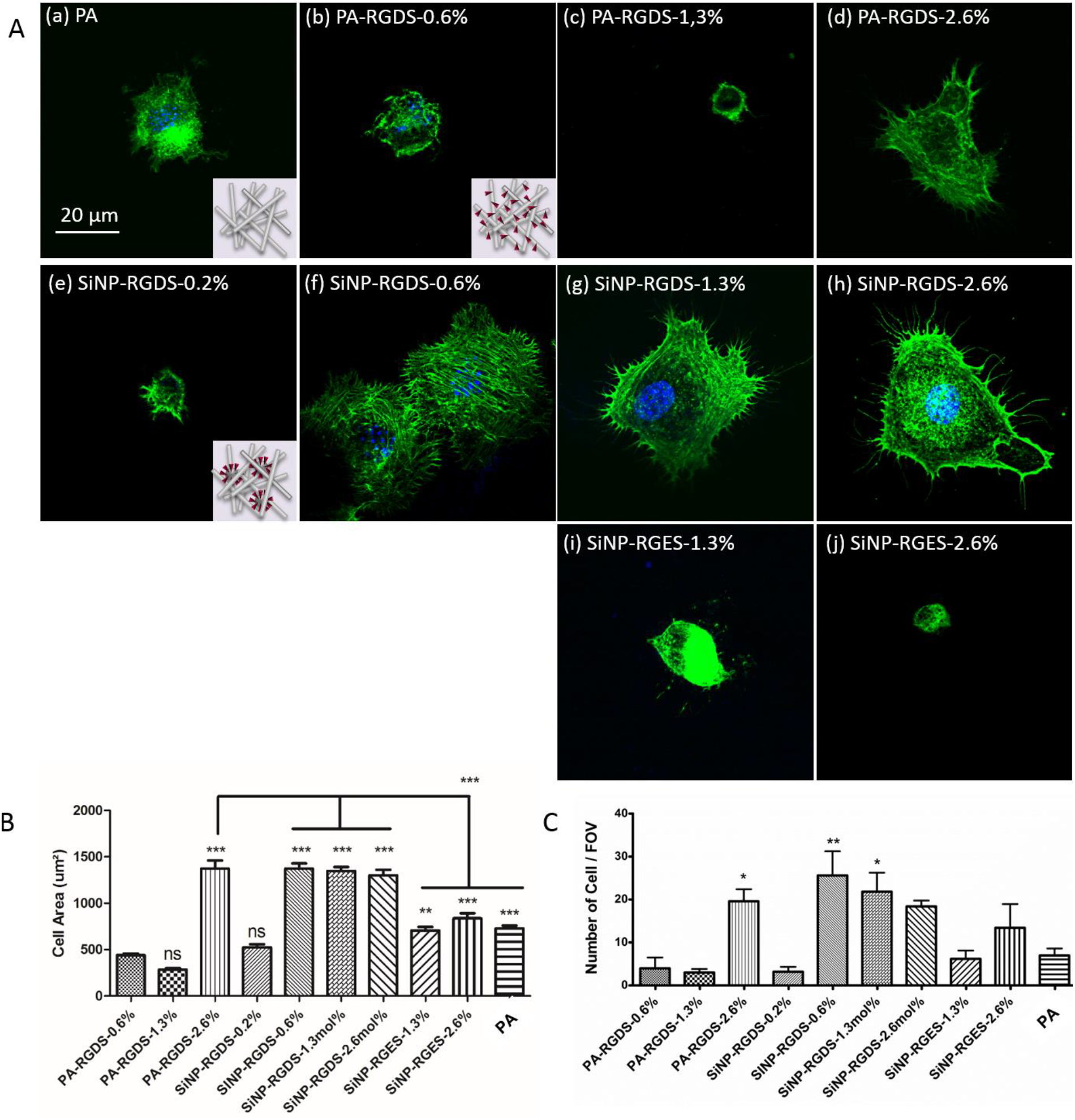
Representative confocal images of 3T3 fibroblasts cultured on PA layers for 4h30 and stained for actin (phalloidin) and nucleus (DAPI) (a) on the PA alone, (b-d) on PA-RGDS, (e-h) on the SiNP-RGDS/PA, and (i,j) on SiNP-RGES/PA negative control at different peptide concentrations. (B,C) Cell morphologies on the different PA layers are compared by measuring the projected cell area and the number of cells by Field of View (FOV). In the plots, the column represents Mean with SEM. (* p < 0.05, ** p < 0.001, *** p < 0.0001; calculated against PA-RGDS-0.6 mol %, unless indicated, using Turkey’s Multiple Comparison test; each condition from three independent experiments).

These results indicate that the clustering of the RGDS bioactive epitopes on the SiNP surface offers an efficient strategy to improve fibroblast cell adhesion on PA matrices.

### 3.3. Bioactivity of divalent-peptide composites: multivalent clustering

The versatility of PA and SiNP chemistry and easy surface functionalization open the possibility of grafting multiple peptide epitopes on the PA and particle surface to promote synergistic binding. This is particularly relevant for mimicking the distance-dependent interaction of the two integrin-binding sequences RGDS and PHSRN. **Figure 5A** illustrates the different possibilities of displaying RGDS and PHSRN peptide epitopes and the modularity of SiNP-PA composites: (1) two separate bioactive PAs bearing RGDS and PHSRN peptides can be co-assembled (PA-RGDS + PA-PHSRN), (2-3) Epitope-modified SiNPs incorporated within a PA matrix modified with the other peptide, *i.e*. SiNP-RGDS in PA-PHSRN scaffold (SiNP-RGDS + PA-PHSRN) or SiNP-PHSRN in PA-RGDS scaffold (SiNP-PHSRN + PA-RGDS), (4) SiNPs grafted with either RGDS or PHSRN incorporated within a peptide-free PA scaffold (SiNP-RGDS + SiNP-PHSRN) and (5) SiNPs grafted with both RGDS and PHSRN within a peptide-free PA scaffold (SiNP-RGDS-PHSRN). For comparison with the previous single peptide (RGDS) experiments, we set the total peptide concentration at 2.6 mol%. Since two peptide epitopes are now used, each was conjugated at a concentration of 1.3 mol% (1.3 mol% RGDS + 1.3 mol% PHSRN). Fibroblast adhesion and spreading on these composites were compared.

**Figure 5.**
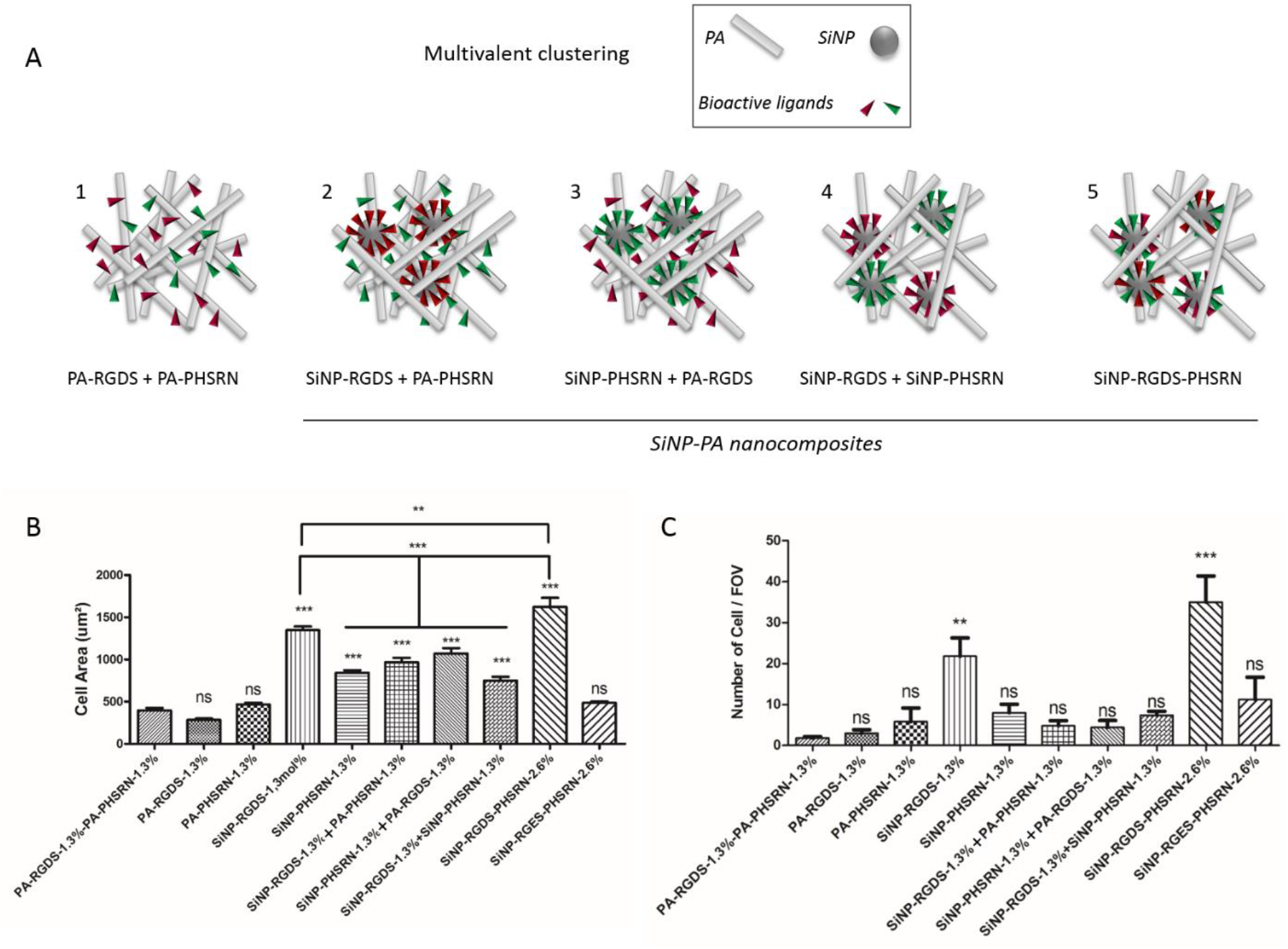
(A) Schematic representation of the different composites prepared to provide multivalent clustering. (1) Bioactive ligands are displayed on PA nanofibers; (2-3) one ligand is displayed on PA nanofibers and the other one is clustered at the surface of SiNPs; (4) the two bioactive ligands are clustered on two different populations of SiNPs; (5) the two bioactive ligands are simultaneously clustered at the surface of a single population of SiNPs. (B-C) Cell morphologies on PA layers are compared by measuring the projected cell area and the number of cells by Field of View (FOV). In the plots, the column represent Mean with SEM. (* p < 0,001, ** p < 0.001, *** p < 0.0001; calculated against PA-RGDS-1.3 mol% + PA-PHSRN-1.3 mol%, unless indicated, using Turkey’s Multiple Comparison test; each condition from three independent experiments).

Quantitative analyses showed limited spreading of 3T3 fibroblasts on monofunctional PA gels (PA-RGDS-1.3 mol% and PA-PHSRN-1.3 mol%) and on matrices obtained by co-assembling the two functional PAs (PA-RGDS-1.3 mol% + PA-PHSRN-1.3 mol%) (**Figure 5B-C**). In contrast, all SiNP-PA composite systems that incorporated the two peptides distributed in the two different phases (SiNP-RGDS/PA-PHSRN and SiNP-PHSRN/PA-RGDS) or on two distinct populations of silica particles (SiNP-RGDS + SiNP-PHSRN PA), promote cell spreading. Interestingly, the composite matrix containing bi-functional particles (SiNP-RGDS-PHSRN) promoted the most spreading. This strongly suggests that the peptide epitopes grafted on the silica nanoparticles are in optimal clusters as well as inter-epitope distance to allow for their synergistic effect on cells.

## 4. DISCUSSION AND CONCLUSION

In this work we integrated silica nanoparticles into peptide amphiphile fibrous networks to yield composite hydrogels. The ability to modify the surface of the particles with bioactive ligands independently from the peptide networks allows for a modular approach to introduce biological cues and control their local density. We demonstrated that surface modification of SiNPs with a diameter of 200 nm ensures the formation of effective peptide clusters with a statistical inter-ligand spacing that can effectively promote cell adhesion and spreading.^4–7^

Furthermore, chemically-engineering the particles to simultaneously display two synergistic bioactive peptides enabled enhanced cell adhesion and spreading. Varying the concentration of surface ligands allowed the control of their density. which was calculated (Figure S5) to be 0.2 molecules per nm^2^ SiNP, *i.e* 1 peptide every 5 nm^2^. This corresponds to a distance of *ca*. 5 nm between the two peptides RGDS and PHSRN, mimicking their separation distance in native fibronectin. It is important to point out that inter-ligand distances can also be controlled by the selected particle size, the synthesis conditions including solvent, surfactant and silica precursor species to vary silanol surface density.^65^ Further control can be achieved by localizing the distribution of peptide epitopes into functional domains or patches.^66^

Major progress in the field of regenerative medicine can be expected from the design of artificial scaffolds that mimic the various features of the ECM. While significant advances have already been achieved in reproducing the structural and mechanical features of the ECM, fine control over the incorporation and positioning of multiple biological cues that play a key role on regulating cell behavior remains highly challenging. This work shows that combining organic and inorganic building blocks with easy, versatile and orthogonal bioconjugation chemistries provides a highly modular approach to engineer 3D scaffolds displaying multiple epitopes. Silica nanoparticles allow for single epitope clustering and multivalent clustering. The possibility to tune their diameters and surface chemistry make them versatile platforms that may be engineered to display multiple epitopes. Integrating bioactive silica nanoparticles with the powerful peptide-based self-assembled matrices generates a new class of composite scaffolds with fine control over the spatial organization of multiple biological epitopes.

## ASSOCIATED CONTENT

## Supporting Information

Characterization of the synthesized peptides; Characterization of the synthesized PAs; SiNPs characterization; Synthesis and characterization of peptide-conjugated SiNPs; Quantification of surface functionalization - Copper-free Click Chemistry Cy3-Azide; Characterization of the self-assembly of the different PA mixtures by TEM; SEM characterization of the PA layers; 3T3 fibroblasts cultured on non-functionalized SiNP/PA composites; 3T3 fibroblasts cultured on PA-PHSRN and SiNP-PHSRN/PA scaffolds.

## AUTHOR INFORMATION

## Corresponding Author

*

## Author Contributions

The manuscript was written through contributions of all authors. All authors have given approval to the final version of the manuscript. ‡These authors contributed equally.

## ACKNOWLEDGMENTS

Peptide synthesis was performed at the Peptide Synthesis Core Facility of the Simpson Querrey Institute at Northwestern University. This facility has current support from the Soft and Hybrid Nanotechnology Experimental (SHyNE) Resource (NSF ECCS – 1542205). Imaging work was performed at the Northwestern University Center for Advanced Microscopy generously supported by NCI CCSG P30 CA060553 awarded to the Robert H Lurie Comprehensive Cancer Center. SEM imaging was performed at the EPIC facility (NUANCE Center-Northwestern University), which has received support from the MRSEC program (NSF DMR-1121262) at the Materials Research Center; the Nanoscale Science and Engineering Center (NSF EEC-0647560) at the International Institute for Nanotechnology (IIN); the Keck Foundation; and the State of Illinois, through the IIN. We thank Mark Trosper McClendon for SEM observations.

## ABBREVIATIONS

PA: peptide amphiphile
SiNP: silica nanoparticle
TEM: transmission electron microscopy
SEM: scanning electron microscopy

